# *Wolbachia* inhibits ovarian formation and increases blood feeding rate in female *Aedes aegypti*

**DOI:** 10.1101/2022.04.19.488749

**Authors:** Meng-Jia Lau, Perran A. Ross, Nancy M. Endersby-Harshman, Qiong Yang, Ary A. Hoffmann

## Abstract

*Wolbachia*, a gram-negative endosymbiotic bacterium widespread in arthropods, is well-known for changing the reproduction of its host in ways that increase its rate of spread, but there are also costs to hosts that can reduce this. Here we investigated a novel reproductive alteration of *Wolbachia w*AlbB on its mosquito host *Aedes aegypti*, based on studies of mosquito life history traits, ovarian dissections and gene expression assays. We found that an extended period of the larval stage as well as the egg stage, as previously shown, can increase the proportion of *Wolbachia*-infected females that become infertile; the impact of *Wolbachia* on infertility therefore accumulated before pupation. We found that ovarian formation was blocked in infertile females, even though infected females had relatively lower *Wolbachia* densities than fertile infected females. Infertile females also showed a higher frequency of blood feeding following a prior blood meal, indicating that they do not enter a complete gonotrophic cycle. Treatments leading to infertility decreased expression of genes related to reproduction. Our results demonstrate effects related to the causes and consequences of infertile *w*AlbB-infected *Ae. aegypti* females with implications for *Wolbachia* releases and have evolutionary implications for *Wolbachia* infections in novel hosts.

## Introduction

*Wolbachia*, a *Rickettsia*-like maternally inherited endosymbiotic bacterium widespread in insects [1], can alter insect reproduction in multiple ways to enhance its transmission [2]. Specific alterations include parthenogenesis – infected females can produce female offspring without mating; feminization – infected genetic males are transformed into functional females and produce female offspring; male killing – infected immature males die while females survive, and cytoplasmic incompatibility (CI) – females infected with the predominant *Wolbachia* strain obtain a fertility advantage because males infected with *Wolbachia* can cause female embryonic lethality when they mate with uninfected females or females infected with a different *Wolbachia* strain [3, 4]. Aside from these alterations that enhance the spread of *Wolbachia*, there can also be costs to the infection that slow its spread [5–7]. These effects of *Wolbachia* are normally based on investigations of natural *Wolbachia* strains that have existed in their hosts for many years, but are also frequently found in artificial infections [8].

In the last decade, artificially introduced *Wolbachia* have been used to reduce the transmission of arboviral diseases, including the *Wolbachia* strains *w*Mel [9] and *w*AlbB [10] in *Aedes aegypti* mosquitoes. Studies on native hosts tend to indicate only a weak fitness cost of *Wolbachia* [6], but *Wolbachia* can have diverse fitness costs in its newly introduced host *Ae. aegypti*, such as life-shortening, heat sensitivity, reduced quiescent egg viability and reduced blood feeding success [11–14]. The effects of *Wolbachia* in native and introduced hosts point to the possibility of a mutualistic relationship between *Wolbachia* developing over time as a consequence of co-evolution and adaptation. In fact, it is acknowledged that sometimes *Wolbachia* can act as both parasite and mutualist [5, 15, 16]. As a result, *Wolbachia* field release and establishment in new populations can be useful for studying evolutionary changes in *Wolbachia* given that the invasion history of the population is known [17–19].

In our previous study of the effects of *Wolbachia w*Mel and *w*AlbB on *Ae. aegypti* eggs, we discovered a novel reproductive effect induced by *w*AlbB, in which infected females that hatched from quiescent eggs lost their fertility despite successful mating, with the proportion of female infertility being correlated with the duration of egg quiescence [20]. The *w*Mel strain also causes female infertility following egg storage, but to a lesser extent than *w*AlbB [20, 21]. Several other reproductive fitness costs of *Wolbachia*, including those on the number and viability of eggs, can be attributed to the consequence of nutritional competition between *Wolbachia* and its host [22] and represent quantitative effects. However, in our study *Wolbachia*-infected females lost their fertility entirely, indicating a qualitative effect difficult to attribute to nutritional competition [20], so we suspect the involvement of other mechanisms.

In this paper, we investigate the causes and consequences of *Ae. aegypti* female infertility induced by *Wolbachia w*AlbB infection after egg quiescence. Firstly, we tested if the effects of *Wolbachia* on female infertility are permanent and dissected female ovaries. Secondly, we performed a larval starvation experiment combined with real-time PCR assays to understand the influence of *Wolbachia* on female development during different stages that could lead to infertility effects. Finally, we tested whether the loss of ovaries in infertile females affected the rate of female blood feeding. Our study highlights the novel nature of reproductive effects altered by *Wolbachia w*AlbB in its new host *Ae. aegypti*, with implications for invasion of *Wolbachia* into uninfected populations and the evolution of mutualism more generally.

## Materials and Methods

### Mosquito populations

We used uninfected and *Wolbachia w*AlbB-infected *Aedes aegypti*. The uninfected population was derived from eggs collected in 2019 from regions in Cairns, Queensland, Australia where *Ae. aegypti* were not infected with *Wolbachia*. The *w*AlbB-infected population was generated by microinjection [23] and infected females were crossed to males from the uninfected population regularly [24, 25] to maintain a similar genetic background between populations. Mosquitoes were maintained in the laboratory following methods described previously [26]. In the following experiments, uninfected females were hatched from eggs stored at 26 ± 1 °C for less than one month. *w*AlbB-infected females from non-stored eggs were hatched from freshly-laid eggs that had been dried and conditioned for one week at 26 ± 1 °C. All lines tested in the same experiment had the same rearing and feeding conditions.

### Determination of female fertility status

To determine the fertility status of individual females in the experiments, we hatched a mixture (approximately 1 : 1) of *w*AlbB-infected eggs that had been stored for two or four months. Female mosquitoes were blood fed on the 4^th^ ± 1 day post emergence. 200 engorged females were aspirated individually in 70 mL specimen cups with larval rearing water and sandpaper strips to allow them to lay eggs. After a week, we separated females that laid (fertile) or did not lay (infertile) eggs and grouped them separately in a 19.7-L BugDorm-1® adult cage (MegaView Science Co., Ltd., Taichung City, Xitun District, Taiwan). The process above is called “fertility separation”. And to test the consistency of female infertility status, we then provided fertile and infertile females with a second blood meal after the “fertility separation”, and isolated 20 of “fertile” and 20 of “infertile” females in 70 mL cups individually for another week to observe if their fertility status was maintained. Next, we dissected the remaining “fertile” and “infertile” females for their ovarian appearance: females were killed by storing at −80 °C for 30 minutes then returning them to room temperature for 10 minutes before dissecting them in saline solution (0.9% sodium chloride) under a compound light microscope (Motic B1 series, Australian Instrument Services Pty. Ltd., Australia).

Since we have confirmed the absence of ovaries from infertile females through dissection, we further improved our method of identifying female fertility status by dissecting females before “fertility separation”. We did this dissection for 65 *w*AlbB-infected females that had been stored as eggs for 11 weeks on the 4^th^ ± 1 day post-emergence (before their first blood meal), and then used an NIS Elements BR imaging microscope (Nikon Instruments, Japan) for photography.

Finally, to test the effect of *w*AlbB on the infertility of *Ae. aegypti* females with different genetic background, we also dissected females from a recent *w*AlbB transinfection line that had been repeatedly backcrossed to a Saudi Arabian background [27]. We dissected ovaries from 113 *w*AlbB-infected mosquitoes that had been stored as eggs for 12 weeks, as well as 50 *w*AlbB-infected females stored as eggs for 1 week and 50 uninfected Saudi Arabian females stored for 12 weeks (Supplementary material 1).

### *Wolbachia* density

After “fertility separation” in the previous section, we measured relative *Wolbachia* density of fertile and infertile females for 16 individuals per group based on the 2^ΔCt^ method [28]. Screening based on real-time PCR assays followed methods described previously [29]. For each replicate, all samples were set up in a 384-well white plate and measured by primers *aeg* and *wMwA* [30]; two consistent replicates (ΔCt < 1) were obtained and their values were averaged for the density analysis.

### Impact of larval stage extension on mosquito infertility

To investigate whether the impact of *Wolbachia* infection on female infertility was only mediated through the egg stage or whether the prepupal stage also had an accumulating impact, we deprived 2^nd^ instar *w*AlbB-infected larvae of food for two weeks before feeding them again until pupation and then followed this with a “fertility separation”. In non-starved controls, *w*AlbB-infected larvae were provided with food *ad libitum*. Each starvation treatment was performed with both stored eggs (stored at 26 ± 1 □ for 10 weeks) and non-stored eggs (stored at 26 ± 1 □ for 1 week) for a total of four treatments. For the group in which infected females were neither stored nor starved, we had two replicates, while for the other three experimental groups we had three replicates, with each replicate containing 30 individual females. Before the blood meal, 15 individual females from each treatment 4 ± 1 days post-emergence were screened for *Wolbachia* infection and *Wolbachia* density, using the screening methods described in the previous section. After the fertile and infertile females had been separated, females were also dissected to confirm their fertility status by checking for the presence of ovaries. We also performed all four treatments with uninfected *Ae. aegypti* (with similar egg storage and larval starvation conditions) and dissected 15 individuals 4 ± 1 days post-emergence from each replicate to check for presence of ovaries (Supplementary material 1).

### Gene expression assays

We selected three genes related to mosquito reproduction and tested their expression levels in females at different developmental stages through a real-time PCR assay [29, 30]. One of the genes tested was the *vitellogenin receptor* (*vgr*): following blood feeding, vitellogenin is secreted from the fat body and internalized by ovaries through the receptor VgR [31–33]. The other two genes were the *ecdysone receptor* (*ecr*) [34, 35] and the *eggshell organizing factor* (*eof*); the vitellogenin transcript is regulated by *ecr* while *eof* is an essential gene encoding a protein for eggshell formation at the late stage of egg production [36] (Supplementary material 2). Gene expression level was quantified relative to a control gene, *RPS17* [37]. We tested female pupae 1 ± 0.5 days post-pupation and female adults 1 ± 0.5 days post-emergence in *w*AlbB-infected females that been stored as eggs for 14 weeks as well as both *w*AlbB-infected and uninfected females that had not been stored, and considered the uninfected mosquitoes as a control. For the *w*AlbB-infected females that had been stored as eggs, we also tested gene expression levels in fertile and infertile females after testing female fertility, and fertile females were treated as the control. Mosquito samples were given a second blood meal and stored in RNA*later*® (Sigma Aldrich Cat No. R0901-100ML) on the third day after feeding, before RNA extraction, reverse transcription and real-time PCR assays were conducted. In all of the above comparisons, we screened 10 individuals from each group; the details of RNA extraction and real-time PCR can be found in Supplementary material 3.

### Blood feeding rate of female mosquitoes

Normally, fully engorged fertile females are reluctant to feed within a gonotrophic cycle. In this experiment, after determining female fertility, we measured blood feeding rate of female mosquitoes by providing fertile and infertile females a third blood meal three days after their second blood feeding, when engorged blood was almost digested. The same volunteer who provided previous blood meals placed their forearm into a BugDorm cage for 15 minutes. Fully engorged females were considered as having successfully fed after this feeding attempt. Three replicates were completed with 20–30 individual females for each replicate. We also weighed 20 random females that had fully fed and 20 that had not been provided with a blood meal from all three replicates to estimate their blood meal weight using a Sartorius Analytical balance BP 210 D (Sartorius, Gottigen, Germany, Readability: 0.01 mg).

### Statistics

We used R v. 3.6.0 with R studio v. 1.1.453 to conduct data analyses and visualizations [38] using the “car” library for ANOVA [39], the base library for other statistical analyses, and the “ggplot2” library for visualization [40]. After real-time PCR measurement, we used the 2^−ΔΔCt^ method to compare gene expression levels [41], using either uninfected females from pupae or young adult stage as controls for the same stage, or fertile females as the control group of infertile females after blood feeding. The values of 2^−ΔΔCt^ were natural log-transformed for ANOVA analysis. For *Wolbachia* density we used the 2^ΔCt^ method [28] to calculate relatively density before log10 transformation for ANOVA analysis. We also used Tukey’s honest significant difference (HSD) tests [42] for further pairwise comparisons. For the proportion data from larval starvation and blood feeding experiments, we analysed data with binomial logistic regression models [43]. In the blood feeding rate test, student’s t-tests were used to test for changes of mosquito weight after blood feeding.

## Results

### Immature ovaries and lower *Wolbachia* densities in infertile *Aedes aegypti* females

We did a “fertility separation” for *w*AlbB-infected females and provided a second blood meal to fertile and infertile females. We confirmed that all females we scored as fertile or infertile after the first gonotrophic cycle maintained their phenotype in the second gonotrophic cycle. We then dissected and compared the ovarian appearance of fertile and infertile females with an Australian or Saudi genetic background and discovered that ovaries could not be observed in infertile females under a microscope (Figure 1). Only occasionally (around one in ten) can immature ovarian structure be seen, but with a narrow width similar to that of a Malpighian tube (Supplementary material 4). We also found that fertile females had higher relative densities of *Wolbachia* (Figure 2: ANOVA: F_1,30_ = 8.632, P = 0.006, fertile: mean ± se = 5.15 ± 0.906; infertile: mean ± se = 2.290 ± 0.673).

**Figure 1.**
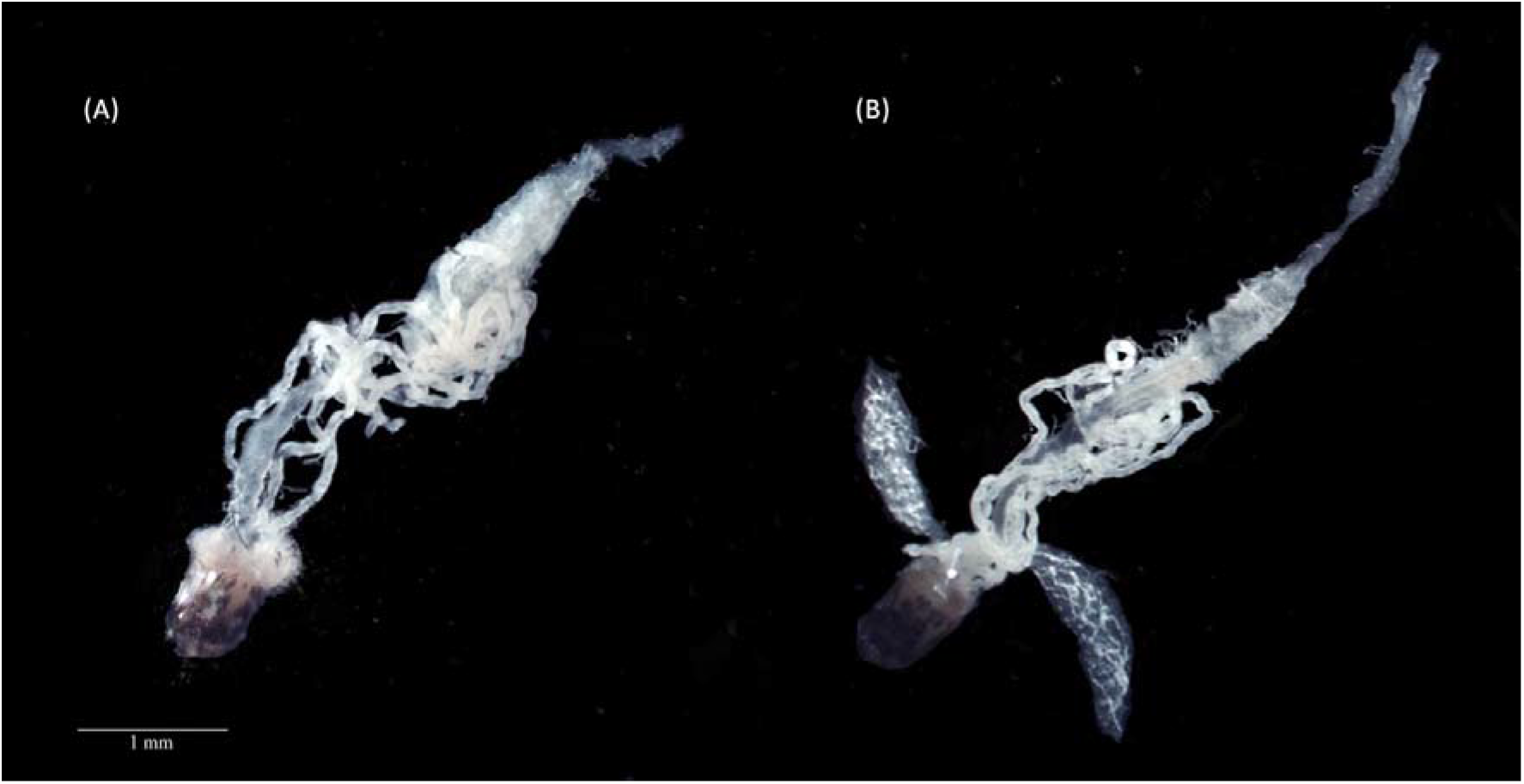
The appearance of internal organs in *w*AlbB-infected *Aedes aegypti* mosquitoes. (a) Infertile females lack observable ovaries, while ovaries are present in (b) fertile females. The background of the photographs was removed for clarity, with the original photographs presented in Supplementary material 4.

**Figure 2.**
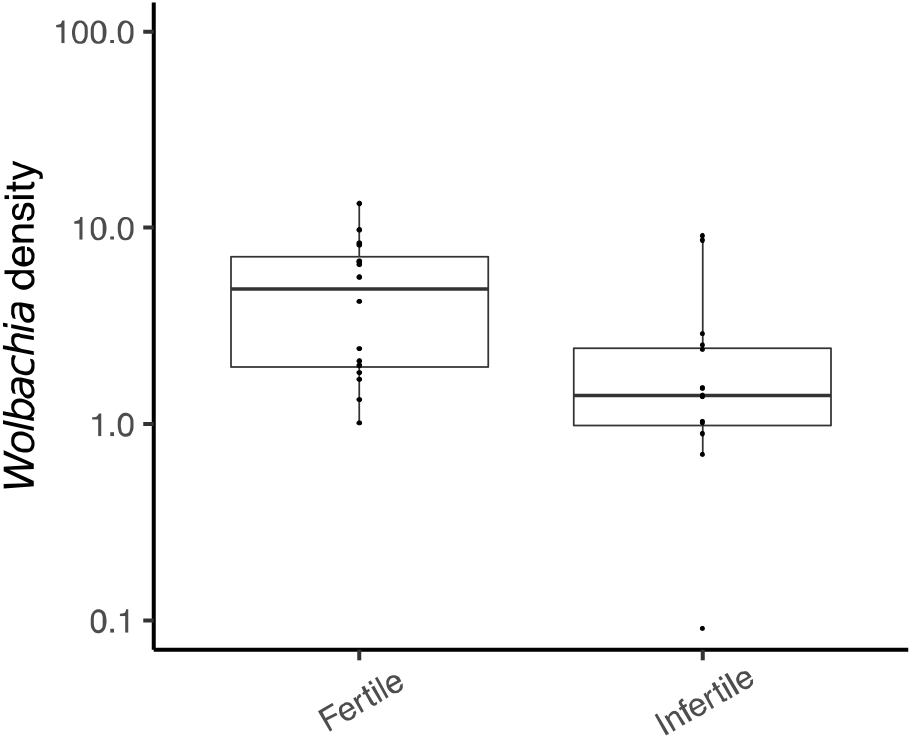
Box plots of relative Wolbachia density of fertile and infertile *w*AlbB-infected females after females were separated into fertile and infertile groups (one week after blood meal). Densities are based on two consistent real-time PCR replicates of 16 females from each group.

### Impact of larval stage extension on mosquito infertility

We starved mosquito larvae to increase the duration of the larval stage from approximately one week to three weeks to test if there is any impact of larval stage extension on infertility in *w*AlbB-infected females. We found significant effects of both storage and starvation treatment, with an interaction between these terms (Figure 3A: storage: F_1,7_ = 256.304, P < 0.001; starvation: F_1,7_ = 32.412, P < 0.001; interaction: F_1,7_ = 21.515, P = 0.002). Infertility rates associated with *w*AlbB infection increased further with extension of the larval stage over and above that seen with stored eggs by 15%, and the extension of the larval stage by starvation also induced infertility of around 10% by itself. For uninfected *Ae. aegypti* treated in the same manner, all females that were dissected contained ovaries, indicating the essential role of *Wolbachia w*AlbB in inducing female infertility during larval starvation and egg storage (Supplementary material 1).

**Figure 3.**
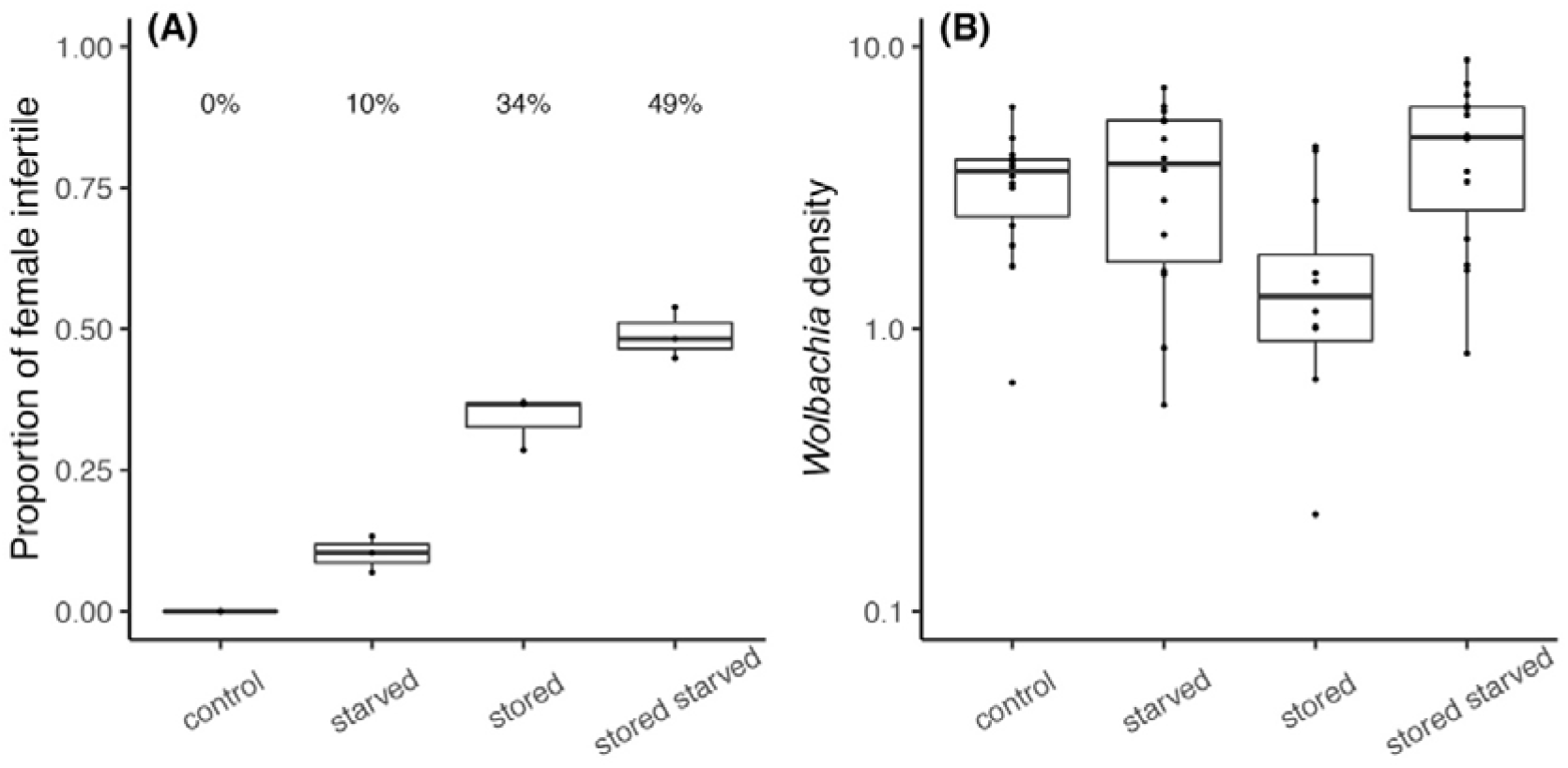
(A) Boxplots showing the proportion of infertile females for *w*AlbB-infected mosquitoes when stored or not stored as eggs and when larvae had been starved or not starved. Values above boxplots represent corresponding averaged percentages based on two replicates for control and three replicates for treatments with each replicate containing 30 individuals. (B) Boxplots of relative *Wolbachia* density in 4 ± 1 days post-emergence females Each group based on two consistent real-time PCR replicates of 15 individuals with failure of detection were excluded (Supplementary material 5).

We also screened for *Wolbachia* in all the treatments to confirm *Wolbachia* infection status. We failed to detect an infection in five out of 60 individuals and these were excluded from the density analysis (Supplementary material 5). *Wolbachia* density was significantly influenced by larval starvation but not egg quiescence, with a significant interaction (Figure 3B: ANOVA: starvation: F_1,51_ = 7.145, P = 0.010; egg quiescence: F_1,51_ = 2.475, P = 0.122, interaction: F_1,51_ = 8.396, P = 0.006). Egg quiescence decreased *Wolbachia* density, but density increased with larval starvation, potentially cancelling out this effect [44].

### Gene expression assays

We selected three genes related to reproductive development and tested their expression level at different developmental stages. At the pupal stage, there are significant differences among the three groups of mosquitoes, but no interaction with the genes (Figure 4A, ANOVA: mosquito group: F_2,81_ = 26.717, p < 0.001; genes: F_2,81_ = 1.044, p = 0.357; interaction: F_4,81_ = 0.423, p = 0.792). Specifically, in Tukey’s HSD test, *w*AlbB-infected females that been stored as eggs had lower expression levels for all three genes than expression levels in uninfected females, but only gene *ecr* had a lower expression level in *w*AlbB-infected females that had not been stored.

**Figure 4.**
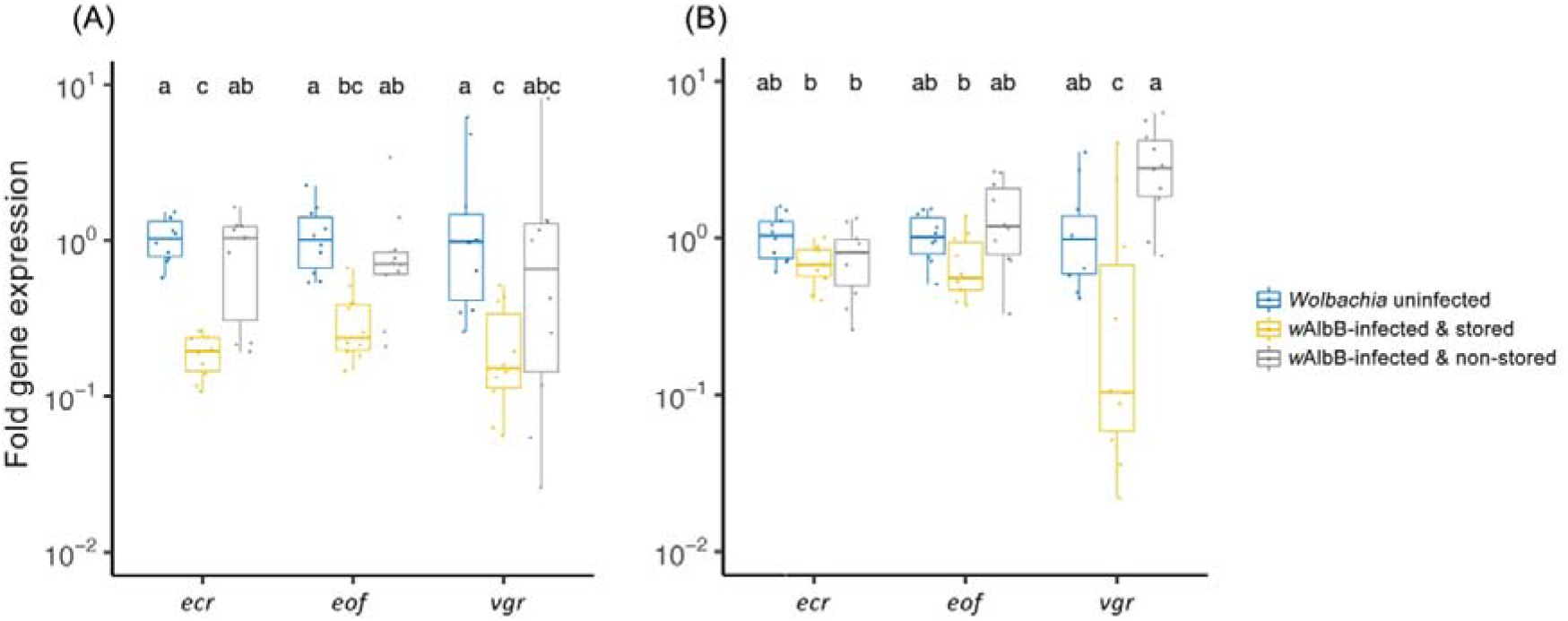
Boxplots of relative expression of reproduction-related genes *ecdysone receptor* (*ecr*), *eggshell organizing factor* (*eof*) and *vitellogenin receptor* (*vgr*) in female (A) pupae and (B) young adults (1 ± 0.5 days post-emergence). Values with the same letter are not significantly different based on Tukey’s HSD tests. Results are based on two consistent real-time PCR replicates of ten individual females from each group.

For young adults 1 ± 0.5 days post-emergence, significant differences of gene expression levels were found among mosquito groups, but there was no interaction with the genes (Figure 4B, mosquito group: F_2,81_ = 15.641, p < 0.001; genes: F_2,81_ = 0.383, p = 0.683; interaction: F_4,81_ = 7.100, p = 0.792). Specifically, only differences for *vgr* expression were significant between *w*AlbB-infected females that been stored and the other two mosquito groups, with some individuals from stored eggs having very low expression levels - these females are probably infertile. The *w*AlbB infection may have increased expression of this gene, indicating that there may also be some impact of *Wolbachia* on reproduction of non-stored females (Figure 4B).

After “fertility separation”, female mosquitoes were provided a second blood meal and were screened for gene expression levels three days later. Significant differences were found between fertile and infertile females, and also between genes and their interaction (Figure 5, mosquito group: F_1,54_ = 235.696, p < 0.001; genes: F_2,54_ = 68.553, p < 0.001; interaction: F_2,54_ = 68.553, p < 0.001). All three genes showed significant differences between fertile and infertile females for expression, especially in the case of *vgr* whose expression in infertile mosquitoes was only around 0.01% of that seen in fertile mosquitoes, likely reflecting the fact that infertile females were unable to produce eggs.

**Figure 5.**
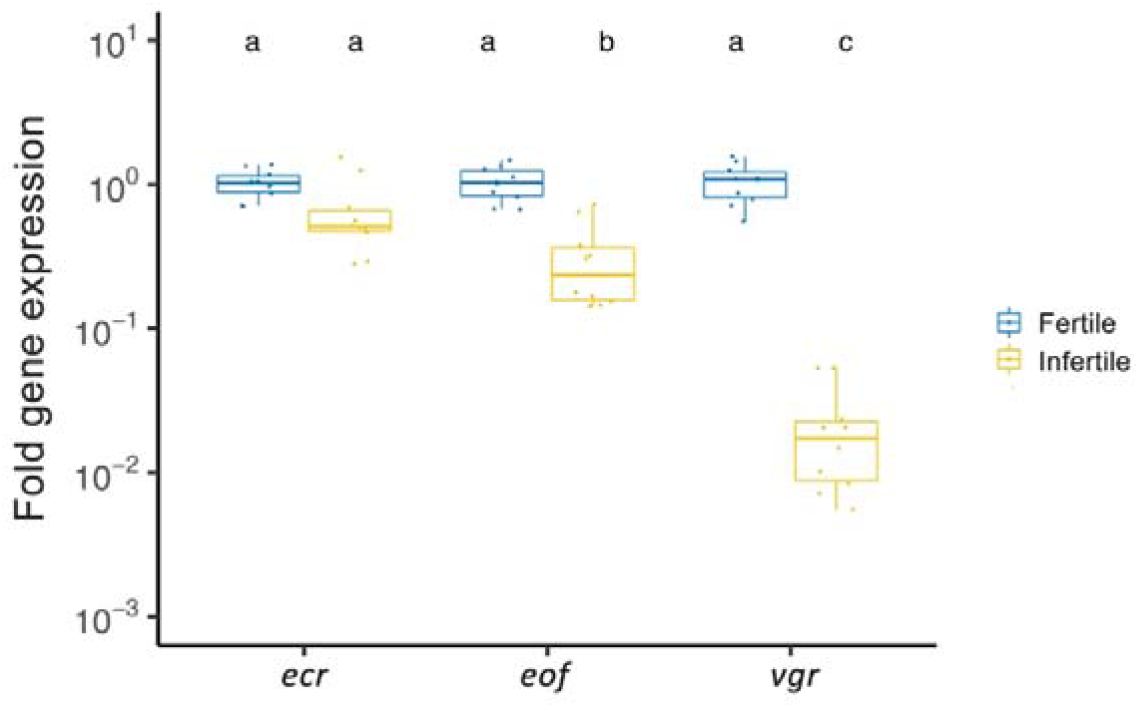
Boxplots of relative expression of reproduction-related genes *ecdysone receptor* (*ecr*), *eggshell organizing factor* (*eof*) and *vitellogenin receptor* (*vgr*) in fertile and infertile females one week after their second blood meal. Values with the same letter are not significantly different according to Tukey’s HSD tests. Results are based on two consistent real-time PCR replicates of ten individual females from each group.

### Blood feeding rate

After “fertility separation”, we provided female mosquitoes a third blood meal on the third day after they had become engorged with their second blood meal to test their blood feeding behaviour. Infertile females had a higher proportion feeding compared to fertile and uninfected females (Figure 6A, F_2,6_ = 61.037, p < 0.001). We also compared the weight of fully fed and unfed females and noted significant differences between mosquito groups, feeding status and their interaction (Figure 6B, ANOVA: mosquito group: F_1,73_ = 29.06, p < 0.001; feeding: F_1,73_ = 322.89, p < 0.001; interaction: F_2,73_ = 15.26, p < 0.001). Specifically, significant weight increases were recorded for both *w*AlbB-infected fertile females (t_35_ = 9.627, p < 0.001) and *w*AlbB-infected infertile females (t_38_ = 15.658, p < 0.001), but the weight of fully fed fertile females was significantly lower compared to infertile females (t_35_ = −4.645, p < 0.001), while there was no significant difference between unfed fertile and infertile females (t_38_ = −1.096, p = 0.280), suggesting that infertile females took in a larger amount of blood during successive feeding.

**Figure 6.**
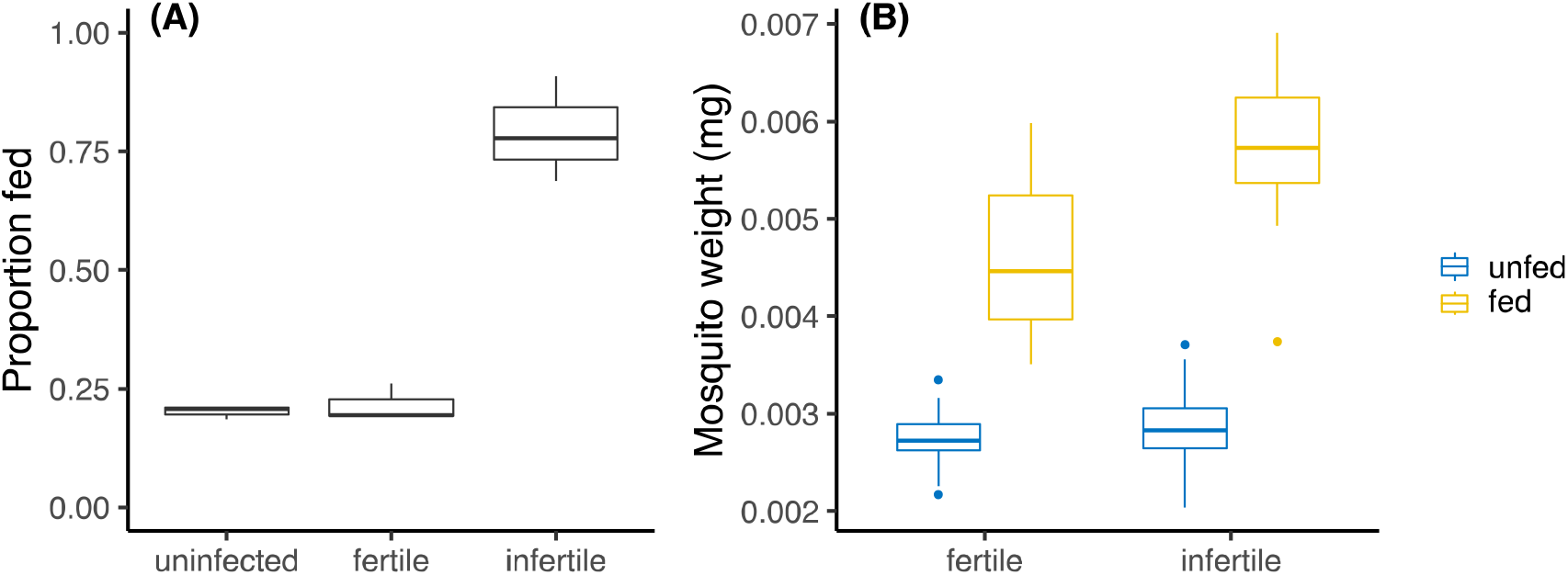
Boxplots of (A) female proportion that blood fed three days after they had been fully fed, in which three replicates were completed with 20–30 individual females for each replicate; and (B) the weights of fed and unfed females. 20 females that had fully fed and 20 that had not been provided with a blood meal from all three replicates were weighed at random.

## Discussion

*Wolbachia* is a well-known bacterium that can alter the reproduction of its host to benefit its spread. In our previous study, *Wolbachia w*AlbB was found to cause *Ae. aegypti* females to become infertile following egg storage [20], inhibiting vertical transmission. In this paper, we investigated this phenomenon and obtained the following results: 1) in infertile females, the development of ovaries has been interrupted by *Wolbachia*; 2) the effect of *Wolbachia* on female fertility accumulates across pre-pupal stages, in that the frequency of female infertility depends on the duration before metamorphosis (including egg and larval stages); and 3) infertile females maintain other female characteristics such as mating and blood feeding, but they do not enter a gonotrophic cycle and blood feeding occurs more frequently. We, therefore, confirmed that *Wolbachia w*AlbB altered the reproductive process in its new host, *Ae. aegypti*.

The effect of *Wolbachia* on female infertility is different from other fitness impacts of *Wolbachia* and the four typical reproductive alterations (parthenogenesis, feminization, male killing and cytoplasmic incompatibility) in which *Wolbachia* alters host reproduction in ways that increase its vertical transmission success [4, 45]. The infertility effect we describe here converts functional females into non-functional females without male characteristics, suggesting different *Wolbachia-related* mechanisms to those involved in other phenomena like feminization [46, 47]. By testing the expression level of *vgr, ecr* and *eof* at the female pupal, young adult before blood feeding and old adult after blood feeding stages, we found substantial differences in gene expression between fertile and infertile groups. For pupae, large variation in expression levels were found among individuals, especially for expression of the gene *vgr*, probably reflecting the fact that its expression mainly occurs in a narrow period during the pupal stage. *w*AlbB-infected *Ae. aegypti* females that had been stored long-term as eggs had lower expression levels of the three reproductive genes compared with uninfected females, especially for the gene *vgr* at the young adult stage, suggesting the formation and development of oocytes was impacted. This is also supported by the very low relative expression of *vgr* and *eof* in infertile females when compared with fertile females at the third day after blood feeding, and the absence of mature ovaries in infertile females. Moreover, we found much lower expression of *ecr* in *w*AlbB-infected female pupae hatched from long-stored eggs, while this difference was not found in young adults. As *ecr* is expressed in a variety of organs to encode the receptor of a hormone [48], the down-regulation of *ecr* may indicate that egg quiescence has a substantial impact on the pupation process of *w*AlbB-infected females. However, interactions between *Wolbachia* and specific reproductive signals and mechanisms are poorly understood and require further investigation for example, through hormone level tests, transcriptome analysis and pathway analysis [49, 50].

The novel reproductive alteration induced by *Wolbachia w*AlbB could slow the spread of the infection in mosquito populations as females are less likely to produce offspring. This may occur not only when the egg desiccation period is extended during dry and warm seasons [51, 52], but also when the larval period is extended due to poor provision of food [53–55]. These effects can reduce the efficiency of *Wolbachia* invasion which aims to inhibit the transmission of arboviruses [8, 25, 56]. While *w*AlbB has established successfully in Malaysia where the warm and humid year-round climate precludes extended egg desiccation periods and promotes fast development of larvae [10, 57], our discovery provides guidance for future releases in other climates and highlights the importance of future monitoring [20].

Our observation that infertile females show an increased rate of blood feeding highlights a potential risk of increased nuisance biting following a *Wolbachia* release program, which may lead to community discontentment. Normal *Ae. aegypti* females ingest human blood to obtain unique nutrients for egg development [58, 59]. However, while infertile females lack ovaries, they still blood feed. Previous research found that multiple feeding within a gonotrophic cycle is caused by nutritional reserve depletion or feeding interruption [60]. Nevertheless, we found a large proportion of infertile females fed again on the third day after their second blood meal, indicating they do not follow a gonotrophic cycle. It is unclear if and how the process of blood digestion is impacted by infertility. Infertile females also took in a larger amount of blood, though this might be a compensatory reaction towards starvation at egg and larval stages. This behavioural change of *Wolbachia*-infected infertile females has significant implications for vector control population release strategies [8, 61]; in particular it would be interesting to test whether the ability of these infertile females to carry and transmit arbovirus has changed as the density of their *Wolbachia* infection may be lower than that of fertile females [62, 63]. Similar concerns have been raised about increased biting frequencies of irradiated female mosquitoes in sterile insect technique programs [64], however the prevalence of infertile females is likely to be much higher in *Wolbachia*-infected mosquito populations compared to irradiated females that are only released accidentally.

The reproductive alterations of *Wolbachia* discovered in this study have evolutionary implications. So far, our work relates to *Ae. aegypti*, where the *Wolbachia w*AlbB was artificially introduced recently from a close relative of *Ae. aegypti, Aedes albopictus*, in order to reduce the transmission of arboviral diseases [65]. A *Wolbachia* strain may induce similar fitness costs in a new host when compared to its native host, such as in the case of life-shortening caused by *w*MelPop which is expressed in both *Drosophila melanogaster* and *Ae. aegypti* [11], and remains stably expressed following long-term laboratory culture [66]. We do not yet know if there are similar reproductive effects of *Wolbachia w*AlbB expressed in its native host, *Ae. albopictus*. On the other hand, we have also discovered a smaller loss of female infertility in *w*Mel infected *Ae. aegypti* [20], though the native host of *w*Mel, *D. melanogaster*, does not enter egg quiescence, although there is a reduced fecundity of *w*Mel-infected females under dormancy conditions [67]. Although other *Wolbachia* strains remain to be investigated, it is possible that *Ae. aegypti* has mechanisms underlying ovarian formation that can be interrupted by endosymbionts. However, it is unclear why the frequency of infertile females increases when the density of *Wolbachia* decreases as the result of increased periods of egg storage or larval development time, and why infertile females have lower *Wolbachia* densities than fertile females in the same treatment. There may be cumulative effects of *Wolbachia* not evident from density measures at early stages of development contributing to failure of ovarian formation. One possible hypothesis which requires further investigation is that *Wolbachia* competes for some essential but sparse nutrients with its host *Ae. aegypti* at an early life stage.

In conclusion, we further investigated a novel reproductive alteration of *Wolbachia* that we discovered previously [20]. We confirmed that *Ae. aegypti* females infected with *Wolbachia w*AlbB can become infertile when they are unable to form functional ovaries during metamorphosis, but these females retain other female characteristics, leading to an increased biting frequency. Our study provides significant guidance for future *Wolbachia* releases and has important evolutionary implications for understanding the reproductive alteration of *Wolbachia*, especially in novel hosts.

## Funding

This research was funded by the National Health and Medical Research Council (1132412, 1118640, www.nhmrc.gov.au). The funders had no role in the study design, data collection and analysis, decision to publish, or preparation of the manuscript.

## Acknowledgment

We thank Véronique Paris for her assistance in weighing mosquitoes.

## Notes

### Competing Interest Statement

The authors have declared no competing interest.

### Summary of Updates

Title and manuscript revised

