## Supplementary material 1 for "*Wolbachia* inhibits ovarian formation and increases blood feeding rate in female *Aedes aegypti*"

Summary of the number of mosquitoes dissected before "fertility separation" and their ovarian developmental status.

| Mosquito genetic background | *Wolbachia* infection | Egg storage duration | Larval starvation | Number with ovaries present | Number with ovaries absent | Number with ovaries immature |
|---|---|---|---|---|---|---|
| Australia | *w*AlbB | 11 weeks | No | 29 | 33 | 3 |
| Saudi Arabia | *w*AlbB | 12 weeks | No | 49 | 57 | 7 |
| Saudi Arabia | *w*AlbB | 1 week | No | 49 | 1 | 0 |
| Saudi Arabia | uninfected | 12 weeks | No | 50 | 0 | 0 |
| Australia | uninfected | 2 weeks | No | 15 | 0 | 0 |
| Australia | uninfected | 12 weeks | No | 15 | 0 | 0 |
| Australia | uninfected | 2 weeks | Yes | 15 | 0 | 0 |
| Australia | uninfected | 12 weeks | Yes | 15 | 0 | 0 |
