## Supplementary material 2 for "*Wolbachia* inhibits ovarian formation and increases blood feeding rate in female *Aedes aegypti*"

Primers used to measure the expression of three essential reproductive-related genes in *Aedes aegypti* females.

| Primer name* | Gene ID | Sequence (5'->3') | Amplicon size (bp) | Efficiency | Source |
|---|---|---|---|---|---|
| ecr838_F<br>ecr838_R | Ecdysone receptor<br>5572184 | CGGAAGGAGAAGAAAGCCCA<br>CGGTAGGTGCTGTTCGTTGT | 80 | 83.77% | This study |
| eof696_F<br>eof696_R | Eggshell organizing factor<br>5576092 | TCCGACCTTGAGCAGCAAAT<br>TTGCTTGCTGGGAGTCTGAG | 75 | 81.64% | This study |
| vgr563_F<br>vgr563_R | Vitellogenin receptor<br>5569465 | GCTTCCGTCGGTACAATCCT<br>GCCTGTGCCGAGAATGAGTA | 93 | 86.46% | This study |
| rps17_F<br>rps17_R | Ribosomal Protein S17<br>110680939 | AAGAAGTGGCCATCATTCCA<br>GGTCTCCGGGTCGACTTC | 200 | 89.04% | [1] |

*Primer name end with '_F' represents forward primer, end with '_R' represents reverse primer, the two with the same prefix are a pair. Correlated genes are shown through the names of the primers.

[1] Dzaki N., Ramli K.N., Azlan A., Ishak I.H., Azzam G. 2017 Evaluation of reference genes at different developmental stages for quantitative real-time PCR in *Aedes aegypti*. *Sci Rep* **7**, 43618. (doi:10.1038/srep43618).
