## Supplementary material 3 for "*Wolbachia* inhibits ovarian formation and increases blood feeding rate in female *Aedes aegypti*"

Within two days after storing at -80°C, ten female mosquitoes from each group were extracted individually for total RNA using the Monarch® Total RNA Miniprep kit (New England Biolabs®, Inc.). Then we used Qubit RNA HS Assay Kit and Qubit 2.0 fluorometer to quantify RNA extractions (Thermo Fisher Scientific) before using High Capacity cDNA Reverse Transcription Kit (Applied Biosystems, Thermo Fisher Scientific, Inc.) to obtain cDNAs (complementary DNA) that were stored at -20°C before further measurements. We used the LightCycler 480 system (Roche Applied Science, Indianapolis, IN) to quantify RNA expression level (real-time PCR). cDNA solution was first diluted six times and 2 μL was pipetted into a 384-well white plate (SSIbio®, Scientific Specialities, Inc., Lodi CA, USA, Cat. No. 3430–40) together with reaction mix buffer. First we examined the efficiency of primers (Table 1) using cDNAs from surplus samples. The DNA amplification began with a 10-minute pre-incubation at 95 ˚C (Ramp Rate = 4.8 ˚C/s), followed by 40 cycles of 95 ˚C for 5 seconds (Ramp Rate = 4.8 ˚C/s), 58 ˚C for 15 seconds (Ramp Rate = 2.5 ˚C/s), and 72 ˚C for 30 seconds (Ramp Rate = 4.8 ˚C/s). In the formal screening we obtained two consistent replicates ($\Delta$Ct < 1) and their values were averaged before analysis.
