## Supplementary material 4 for "*Wolbachia* inhibits ovarian formation and increases blood feeding rate in female *Aedes aegypti*"

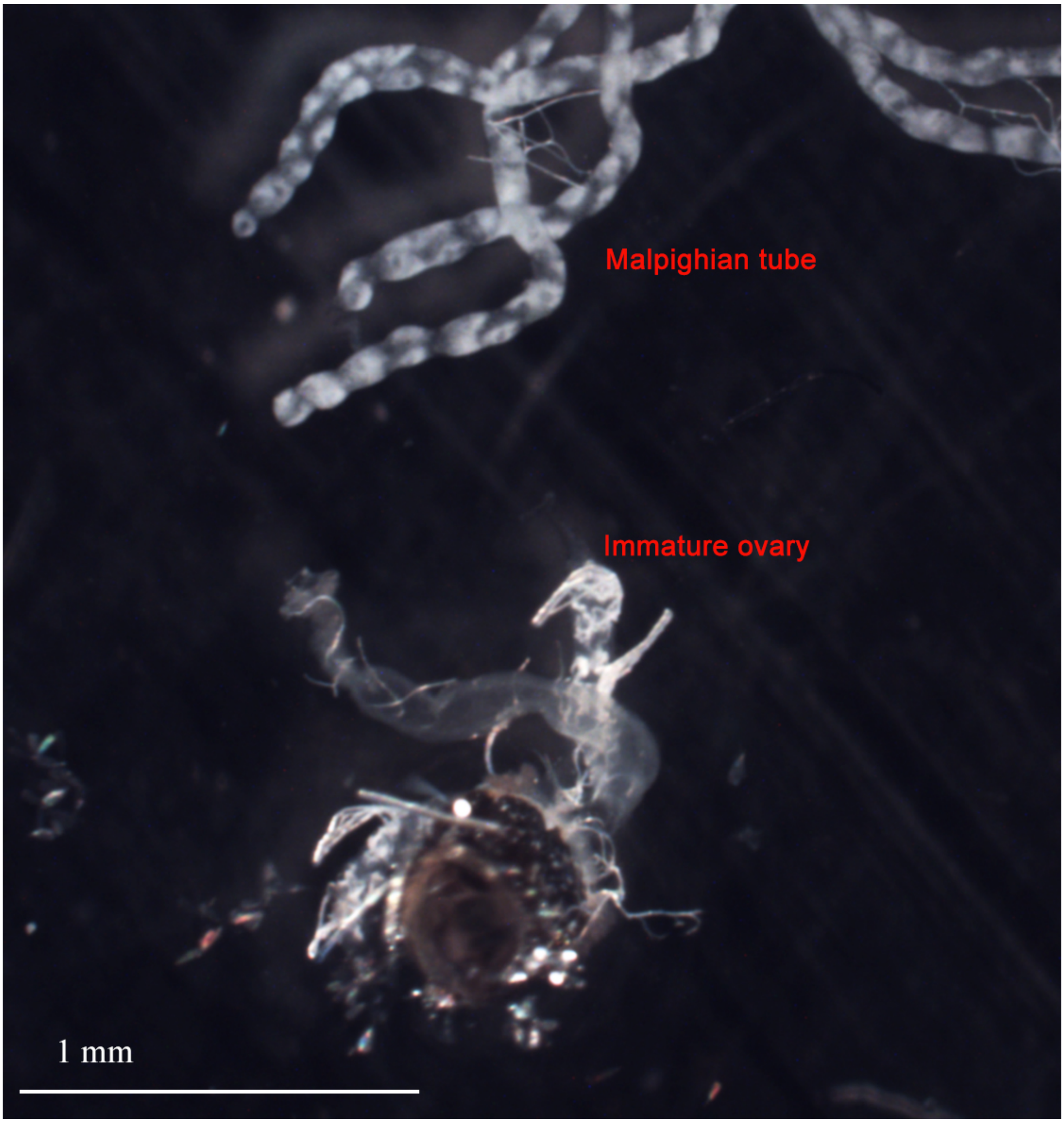


Sup Figure 1. A rare case where immature ovarian structures can be seen in infertile females, with the width of ovaries similar to Malpighian tubules.

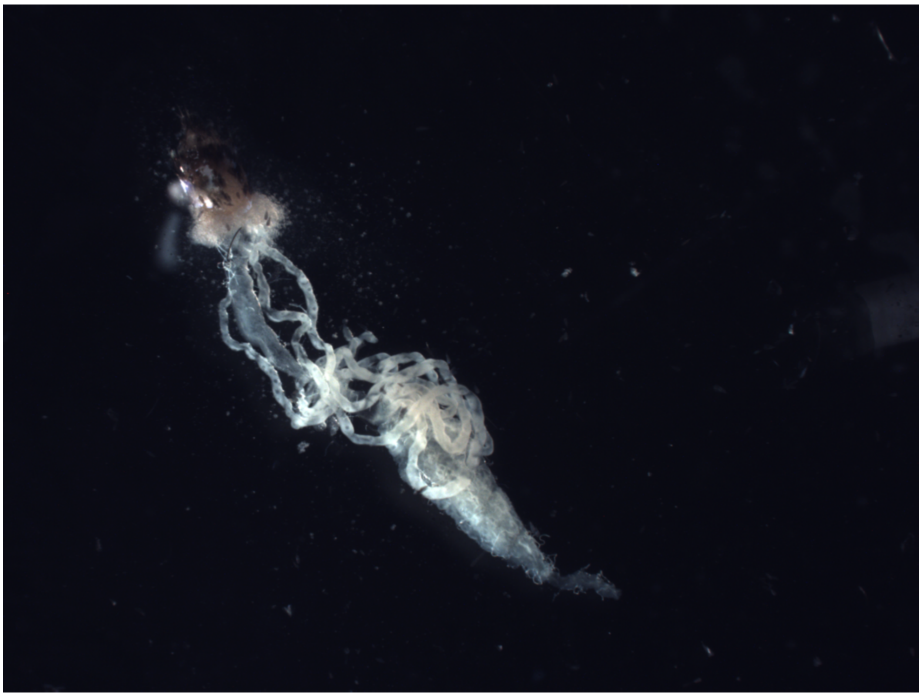

Sup Figure 2. Original picture of Figure 1A.

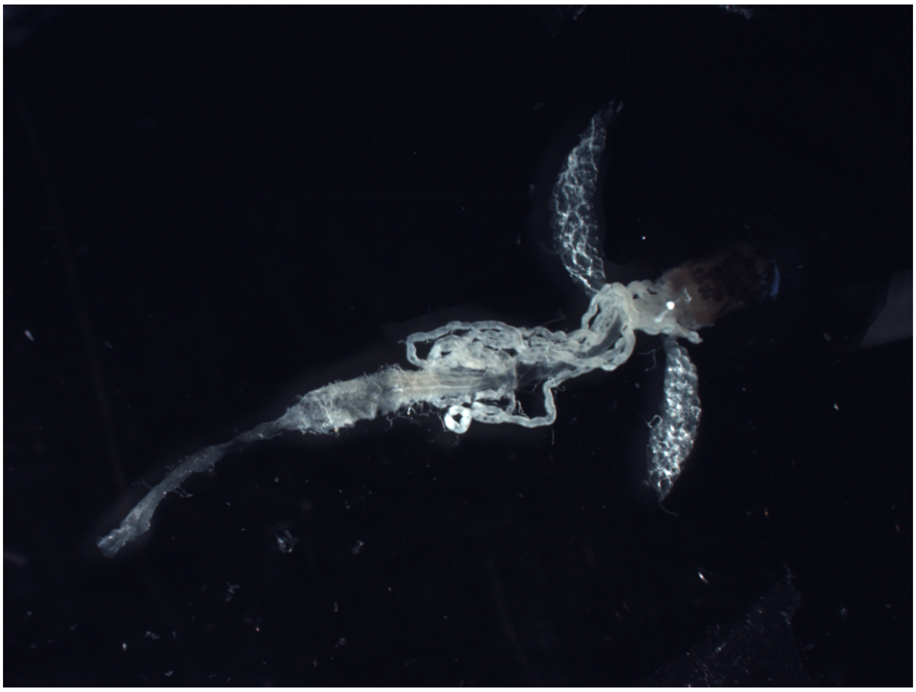

Sup Figure 3. Original picture of Figure 1B.
