## Supplementary material 5 for "*Wolbachia* inhibits ovarian formation and increases blood feeding rate in female *Aedes aegypti*"

Summary of the number of uninfected mosquitoes that were excluded in the density analysis in the *Ae. aegypti* larval starvation experiment.

| Colonies | Storage (Y/N) | Starvation (Y/N) | Number of uninfected |
|---|---|---|---|
| control | N | Y | 1 |
| stored | Y | N | 3 |
| starved | N | Y | 0 |
| stored starved | Y | Y | 1 |
